# Using Mixtures of Biological Samples as Process Controls for RNA-sequencing experiments

**DOI:** 10.1101/015107

**Authors:** Jerod Parsons, Sarah Munro, P. Scott Pine, Jennifer McDaniel, Michele Mehaffey, Marc Salit

## Abstract

**Background:** Genome-scale “-omics” measurements are challenging to benchmark due to the enormous variety of unique biological molecules involved. Mixtures of previously-characterized samples can be used to benchmark repeatability and reproducibility using component proportions as truth for the measurement. We describe and evaluate experiments characterizing the performance of RNA-sequencing (RNA-Seq) measurements, and discuss cases where mixtures can serve as effective process controls.

**Results:** We apply a linear model to total RNA mixture samples in RNA-seq experiments. This model provides a context for performance benchmarking. The parameters of the model fit to experimental results can be evaluated to assess bias and variability of the measurement of a mixture. A linear model describes the behavior of mixture expression measures and provides a context for performance benchmarking. Residuals from fitting the model to experimental data can be used as a metric for evaluating the effect that an individual step in an experimental process has on the linear response function and precision of the underlying measurement while identifying signals affected by interference from other sources. Effective benchmarking requires well-defined mixtures, which for RNA-Seq requires knowledge of the messenger RNA (mRNA) content of the individual total RNA components. We demonstrate and evaluate an experimental method suitable for use in genome-scale process control and lay out a method utilizing spike-in controls to determine mRNA content of total RNA in samples.

**Conclusions:** Genome-scale process controls can be derived from mixtures. These controls relate prior knowledge of individual components to a complex mixture, allowing assessment of measurement performance. The mRNA fraction accounts for differential enrichment of mRNA from varying total RNA samples. Spike-in controls can be utilized to measure this relationship between mRNA content and input total RNA. Our mixture analysis method also enables estimation of the proportions of an unknown mixture, even when component-specific markers are not previously known, whenever pure components are measured alongside the mixture.

## Background

Measurement assurance for genome-scale measurements is challenged by the impracticality of creating a sample containing known quantities of tens of thousands of components, such as the RNA transcripts measured in an RNA-seq experiment. Deep sequencing of cellular RNA can generate vast quantities of gene expression information, yet measurement biases have been identified at nearly every step of the library preparation process [1-4].

As RNA-sequencing expression data expands from discovery into clinical applications, the sources and magnitudes of bias and variability must be carefully understood and quantified. The basic units of expression in sequencing, such as transcripts per million reads (TPM) or fragments per kilobase per million reads (FPKM), are still undergoing revision [5,6]. Even when using comparable units, it is rarely possible to directly compare gene expression values reported by different labs, on different instruments, or frequently just on different days [6-8], unless special care is taken to use uniform samples and protocols. Identifying the presence and variation of biases in a measurement process over time requires a standard to be used for process control. The regular use of a process control can help determine the most-appropriate protocol and analysis methods, demonstrating that they accurately represent the true changes in the underlying sample.

Ideally, a measurement process is linear and possesses a known precision. A linear measurement process shows an increase in signal proportional to an increase in the object being measured. It is also helpful if measured signal is additive, arising only from a single source. Precision consists of repeatability and reproducibility, defined as the degree of closeness in multiple measurements made by a single user and the closeness between multiple labs, respectively. We show that mixtures can demonstrate that a measurement’s response function is linear and of high specificity (free of interference or cross talk) while measuring its variability and precision. Properly constructed mixture samples can be used to correct for systematic measurement errors, provide ongoing monitoring of performance, serve as a tool for interlaboratory comparison, and create a context for evaluating batch effects, protocols, and informatic analyses.

There are two approaches to creating useful genome-scale standards. One is the creation of a limited number of external spike-in controls, such as those designed by the External RNA Controls Consortium (ERCC), which were created for microarrays and have been applied to next-gen sequencing [9-11]. A second approach utilizes mixtures of previously characterized samples in defined ratios, and has also been applied to microarrays [12-14] but has not been utilized in other genome-scale measurements. Using these types of standards provides confidence in the ability of a test to detect both positive and negative results, determining the limits of that detection.

Mixtures can serve as a test that applies to each of the tens of thousands of transcripts in a sample’s transcriptome. Linearity of the measurement response can be demonstrated based on the fundamental understanding that a mixture is a linear combination of its components. Previous work with mixtures in microarrays[12-14] utilized an arbitrary 10-fold “selectivity” cutoff to evaluate the linear dynamic range of microarray measurements and understand the variability of these measurements. The arbitrary selectivity cutoff in previous work prevents the identification of interference, as any genes affected by interference would be filtered by the stringent selectivity cutoff.

Using known mixture compositions, predicted values can be calculated based on the assumption that the measurement response is linear. Deviation of the observed values from the model-predicted value is an indication of bias in the measurement. Systematic biases could be introduced by sample preparation, signal processing, interference from related or mis-annotated genes, or sampling variation. Signal arising from off-target molecules, such as a closely related transcript, can cause false positive results and result in a lowered specificity. Mixture samples can provide information about the measurement sensitivity, specificity, repeatability, reproducibility, dynamic range, and limit of detection.

Determining the relative contributions to gene expression of individual components within mixtures of biological states has received some attention in clinical research, where biopsies and other patient samples are often mixtures. The process of resolving gene expression signals introduced by each individual component of a mixture [13-23] has been used to account for tumor heterogeneity and to separate whole blood samples into individual cell types. These procedures often separate mixture components based on a subset of genes forming a signature that varies uniquely between components. These deconvolution methods have been used [27-30] to develop high-resolution tumor expression signatures from imperfect biological samples [31,32] and differentiate between cell-type-frequency changes and per-cell gene expression changes [33,17]. Many of these methods can determine mixture component types by using a linear model where mixture expression is treated as a combination of expression signatures.

One parameter notably absent from these methods is RNA content. Different cell types express different total amounts of RNA per cell, confounding estimates of cell type proportion made based on the quantification of total RNA [24]. Others have introduced the concept of a biological scaling factor [25,26] to compensate for variation in the RNA content of cells, including the use of spike-in controls to determine this factor. The enrichment of mRNA from total RNA adds a bias to the experiment due to the different abundance of mRNA between cell types.

We assess linear response, specificity, and accuracy of genome-scale measurements using mixtures. In the process, we demonstrate that linear models can be used to separate these mixtures into the proper components. We were mindful that while our mixtures were of total RNA, the sequencing process filters for mRNA, and that the relationship between these two values is an important factor when interpreting results. We anticipate that a mixture-based approach to measurement assurance is highly generalizable to many types of mixtures and can be extended to the wide variety of genome-scale measurements, including but not limited to proteomic and metabolomic measurements.

## Results

To assess measurement parameters of genome-scale transcriptome data, we analyzed two RNA-seq experiments measuring synthetic mixtures of commercially available human total RNA samples (Figure 1)[13,14,34]. One experiment included a mixture of two reference total RNA samples, sequenced by 9 laboratories as a part of the Sequencing Quality Control Consortium (SEQC) [34-35]. In this study, the 9 laboratories sequenced the following samples: Universal Human Reference RNA spiked with ERCC ExFold RNA Spike-in Mix 1 (SEQC-A), Human Brain Reference RNA spiked with ERCC ExFold RNA Spike-in Mix 2 (SEQC-B) and two mixtures of SEQC-A and SEQC-B (SEQC-C and SEQC-D) with mixture compositions C=3A+1B and D=1A+3B. These four samples were sequenced by 9 labs using either Illumina or Life Technologies sequencing instruments.

**Figure 1:**
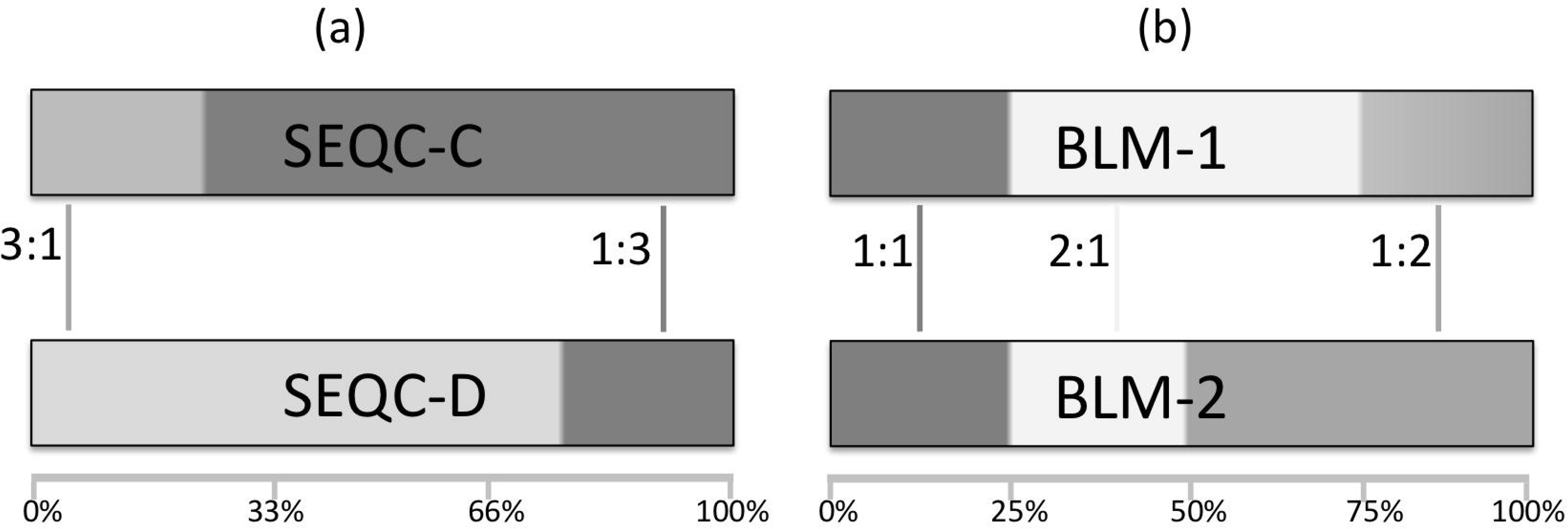
RNA samples used in this study. RNA isolated from pure tissues is used to generate pairs of mixtures used in two separate experiments. (a): Two SEQC mixtures (SEQC-C and SEQC-D) are built from two components (SEQC-A and SEQC-B). (b): Two BLM mixtures (BLM-1 and BLM-2) are built from three components. The SEQC-B component (HBRR) is from the same source as the Brain BLM component. Per-sample target ratios of tissue proportion between mixtures are shown.

The second sample, BLM, contains two mixtures (BLM-1 and BLM-2) composed of total RNA isolated from human brain (B, the same RNA as SEQC-B), liver (L), and muscle (M) tissue. These two mixtures were made with component proportions of 1B:1L:2M and 1B:2L:1M. The total RNA of each individual tissue were also sequenced as single component samples to provide an expression signature for each tissue. ERCC spike-in control RNAs[12] prepared by NIST were added to the BLM mixtures and individual components. Two spike-in control pools were designed with ratiometric differences in the concentration of individual ERCC spike-ins. As expected based on the mixture designs, ERCCs spiked-in equally yielded equal expression signal, while signal from ERCCs spiked differentially into multiple subpools was at ratios corresponding to the designed fold changes. Poisson sampling at the lower expression levels results in increased dispersion about the expected ratio [49].

These mixtures were designed to have a defined expression signal ratio between them. For example, if the measurement response were linear and unbiased, the signal in the SEQC-C sample would be exactly 1/4 the signal of SEQC-B plus 3/4 the signal from SEQC-A due to the design of the mixture. However, these total RNA mixtures went through RNA-seq library preparation, which purposely removes ribosomal RNA from the sample. The resulting sequence data reflects this filtration, which can be different from sample to sample. A correction for this differential enrichment, including upper-quartile normalization[36], must be applied to accurately reflect the experimental process and allow the model to return the designed ratios of expression between mixtures (Supp.Figure 1).

### Linear model-based analysis of genome-scale gene expression

We observed that mixture expression is a linear combination of the component samples and the mixture proportions of each component. Equation 1 describes the relationship between signal in the mixtures and signal in the constituent samples. A mixture *M* (two per dataset in this study) is composed of a number of named components *C* (“B”,”L”, and “M” in the Brain/Liver/Muscle mixture or “A” and “B” in the SEQC dataset), with each component comprising a proportion of the mixture *Φ*_C_. *χ*_i,M_ is the expression signal arising from a particular gene/transcript *i* in mixture *M*.

**Equation 1**: 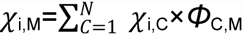

This study uses four mixtures of the same general form:

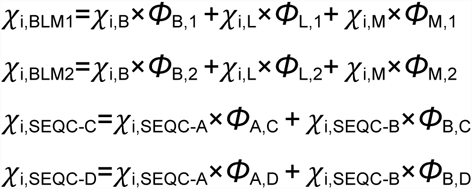

These mixtures were made from total RNA, while the expression signal (sequencing reads) arises only from the mRNA. As the fraction of the total RNA mass that is mRNA varies between cell types, the filtering of total RNA into mRNA introduces a bias. Supplemental Figure 1 shows the offset from the expected ratios of tissue-specific and ERCC RNA caused by this bias. We correct the specific equations for the mRNA fraction by multiplying each component by a factor *ρ*. This factor corresponds to the measured mRNA compared to the mass of total RNA in each mixture. *ρ*_*C*_ is defined as the amount of measured RNA per unit total RNA in component *C*.

Including this factor, the BLM1 mixture equation becomes Equation 2:

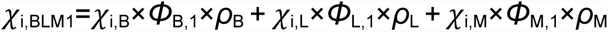

There are a few approaches that have been described to measure *ρ*. One study directly measured the mRNA content between SEQC-A and SEQC-B samples [36]. Another described the use of trimmed mean of log expression ratios (TMM)[25] to measure mRNA content from RNA-seq data. TMM-derived factors have been shown to be an appropriate measure in cases where there is no global expression level change (such as the SEQC mixtures), but introduce bias if there are global expression changes (such as in the BLM mixtures)[26].

The *ρ* factor can be determined using spiked-in RNA[26] as sample reads per microgram of total RNA divided by spike-in reads per microgram of spike-in RNA. This calculation emphasizes that the mRNA fraction is a correction for the differential enrichment between polyadenylated spike-in RNA and total RNA, which is only partly composed of mRNA.

Figure 2 compares the distributions of spike-in estimated rho factor ratios across the SEQC samples compared to the direct measurement of mRNA quantities in total RNA made previously [36]. While the *ρ* factors do not permit direct mRNA content measurement, the ratiometric measurements of pairs of samples have distributions that are similar to that of a normal distribution with parameters based on the previous mRNA content measurements of SEQC-A and SEQC-B. Additionally, the expected equalities of *ρ*_C_= *ρ*_A_*.75+ *ρ*_B_*.25 and *ρ*_D_= *ρ*_A_*.25+ *ρ*_B_*.75 hold true to within 5% of *ρ*_A_, indicating that the mRNA content of a mixture is a linear combination of the mRNA content of its components. Additionally, solving the system of BLM equations only for the mRNA fractions (inputting the known proportion values) yields very similar mRNA fractions to those calculated from spiked-in RNA, leading us to be confident in these measurements.

**Figure 2:**
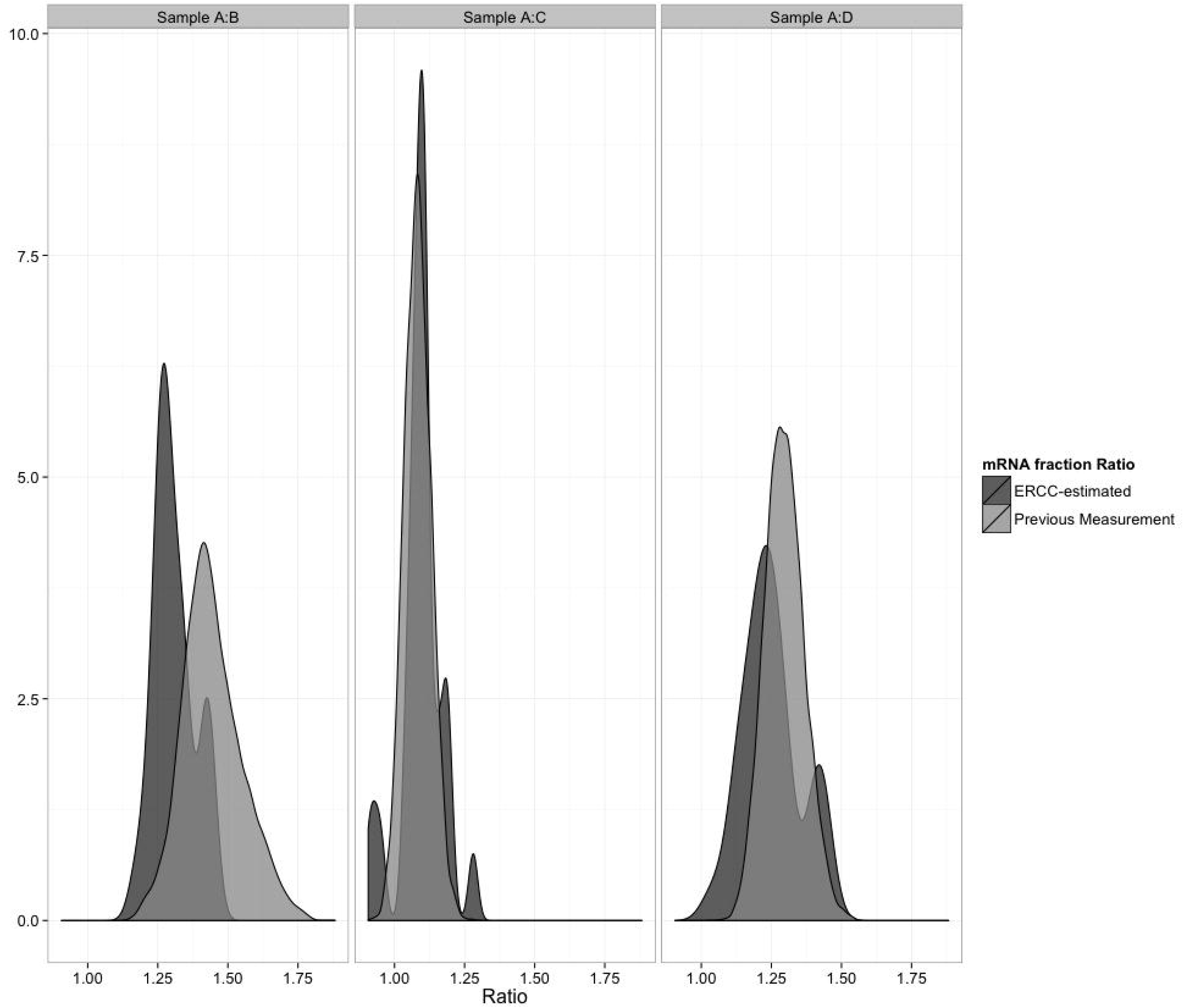
Distributions of empirical (light) and ERCC-estimated (dark) mRNA content ratios between SEQC samples A:B, A:C, and A:D. The empirical distribution was simulated from a normal distribution with means of 2.87 and 2.003 and standard deviations of 0.095 and 0.124 for samples A and B, as reported previously [37]. The ERCC-estimated values were calculated from Equation 3. Individual labs’ mRNA enrichment varied inside a narrow range, yielding discrete peaks in the distribution for some outlying labs.

The mRNA fraction *ρ* is a property of an individual RNA sample and is affected by any RNA manipulation - particularly the mRNA enrichment step in sample preparation. For replicates within a single polyA-selected SEQC experimental run, the *ρ* of a mix varies slightly, likely due to fluctuations in efficiency of mRNA enrichment. (S.Table 1) It is also important to note that FPKM units should not be used to calculate mRNA fraction (Supp.Figure 2), as the FPKM derivation [6] includes a term coupling sample abundance to spike abundance.

### Mixture analysis models recapitulate known mixture proportions

To demonstrate the accuracy of this analytical framework of mixtures, the mixture proportions *Φ*_*BLM*_ were recalculated for the BLM mixtures BLM-1 and BLM-2. The *ρ* values and the sequencing expression data X_i_ were used to solve for the mixture proportions *Φ*_*BLM*_ by linear regression to the mixture equation. Figure 3 shows that the experimentally observed counts are highly correlated (R^2=0.996) to the equation-solved counts X_i_ for each transcript. Figure 4 shows the *Φ*_*BLM*_ values at which residuals were minimized for the two mixtures for each replicate sample in each laboratory. Estimates of the three component proportions in the two mixtures are consistent with the designed 25:25:50 and 25:50:25 proportions in the two BLM mixtures. Figure 5 shows that the designed proportions of SEQC mixtures across each of nine labs can also be calculated by this equation, returning the 75:25 and 25:75 proportions for mixes C and D, with some variability between labs. Equation 1, which lacks correction for mRNA fraction, does not return the designed ratios (Supp. Figure 3).

**Figure 3:**
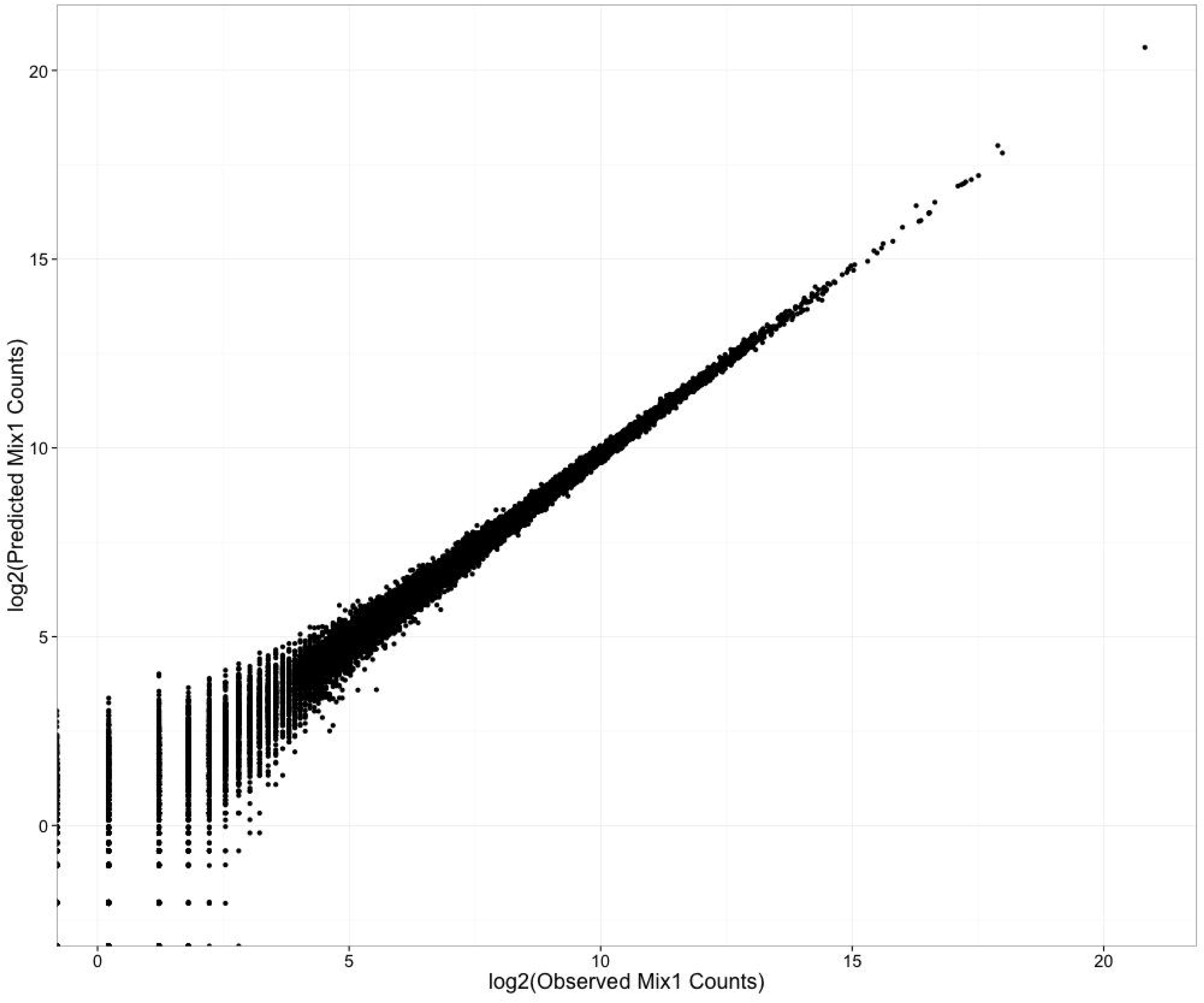
Comparison of observed and predicted counts. Observed BLM mix 1 counts (x) are plotted against predicted BLM mix 1 counts (y). Predicted counts are calculated using equation 2. Counts are on the log2 scale.

**Figure 4:**
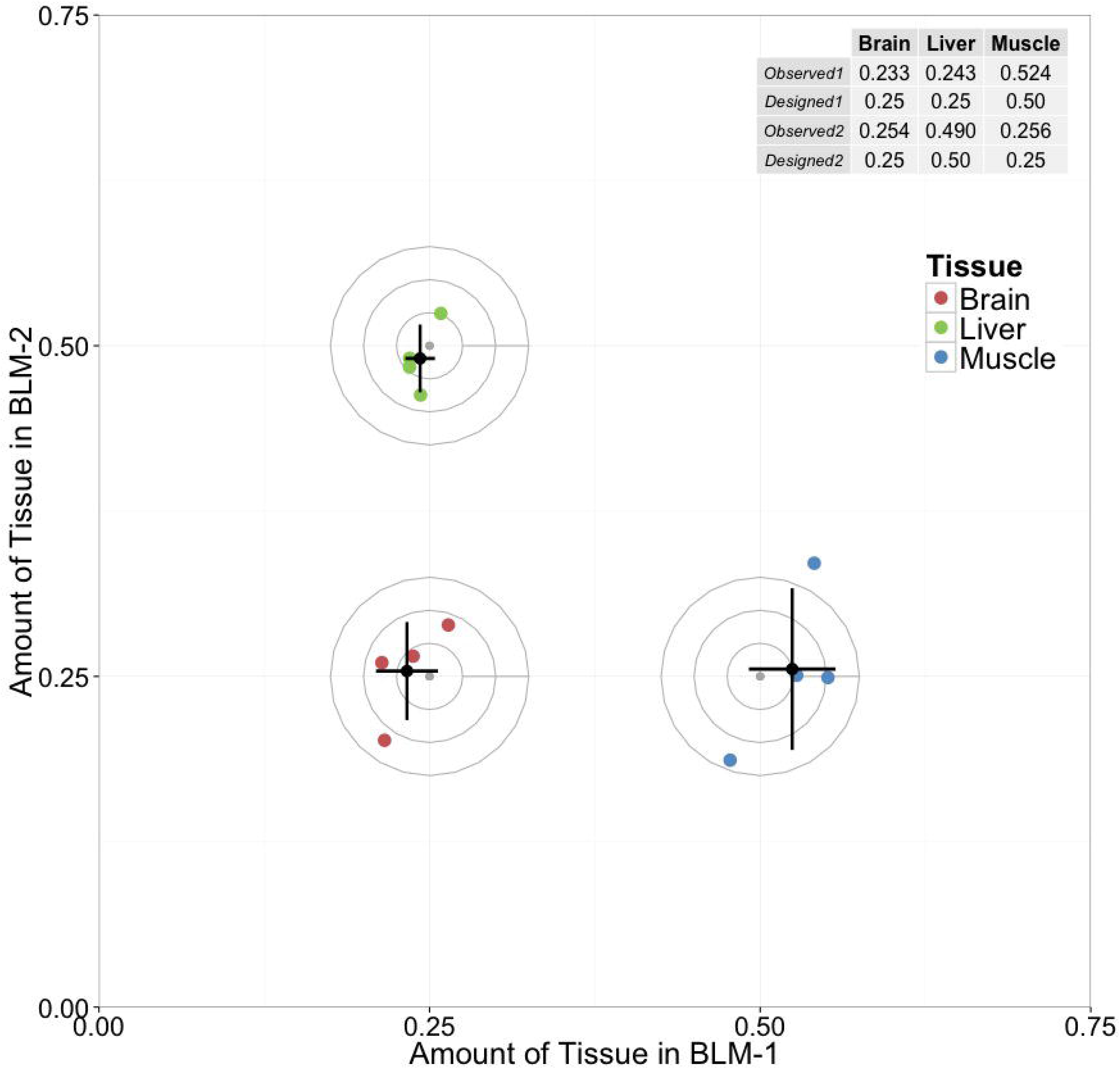
Accuracy of model-derived BLM mix estimates. The grey center point is the nominal ‘truth’ ratio in which the samples were mixed. Concentric circles with radius at multiples of 0.025 are added to visually clarify distance from the center point. Colored points depict mixture proportion *(Φ)* estimates generated from measurements of 4 replicate libraries. Black points are the mean of the replicates. Error bars show one standard deviation of the four replicate measures

**Figure 5:**
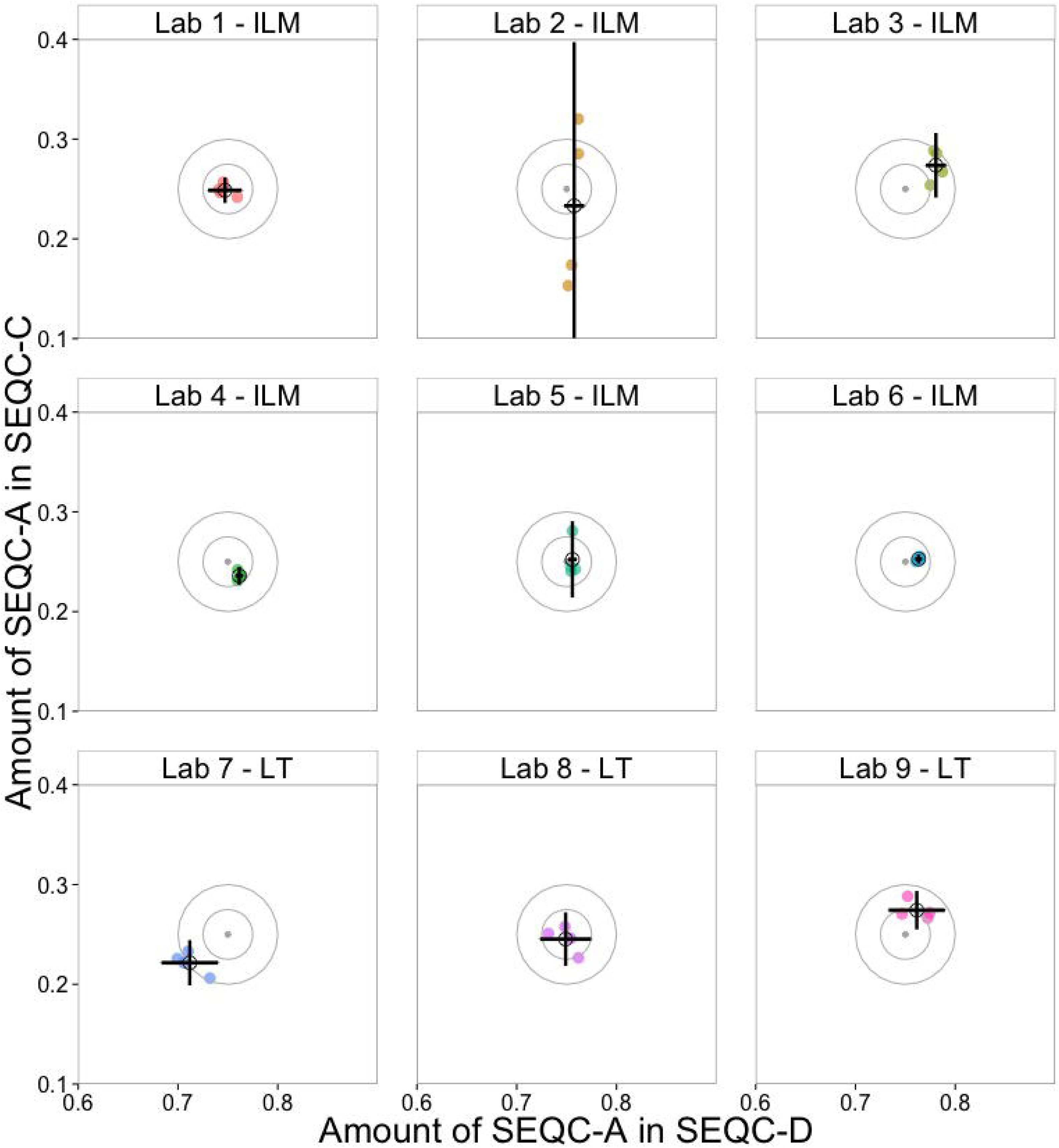
Mixture proportion (*Φ*) estimates for samples A in SEQC-C and SEQC-D. The mean (black hollow circle) and standard deviation (error bars) of four individual replicates (colored) of the *Φ* estimate for each sample are shown. The nominal mixture proportions are grey points at the center of the target. Circles centered at that nominal ratio with radii in multiples of. 025 are included to more easily identify magnitude of total error. LT and ILM tags indicate the manufacturer of the sequencer used at each lab (Life Technologies and Illumina, respectively). Deviations from the target indicate process variability, instrument bias, or errors brought about in these labs.

### Linear model-predicted mixture counts are equivalent to replicate measures

In studies by the SEQC [34], differential expression between replicate samples was utilized to evaluate measurement performance based on the hypothesis that the control samples used in the study had no true differences between replicates. We created pseudo-replicate predicted count values from the single component samples for use in benchmarking. These simulated mixtures were built based on the measured mixture expression and the true mixture proportions.

Figure 6 shows a dendrogram of the distance between actual mixture expression and predicted expression counts of SEQC samples. The four base samples A, B, C, and D are most distant from one another, reflecting the biological differences between the samples. Samples A and C are more closely related, as C consists of 75% A and 25% B. Modeled pseudo-replicate samples Cm and Dm across each of the six SEQC sites are no more different than cross-lab replicates of the C and D data, indicating that building the model for mixture C from components A and B does not introduce significant variability. This supports the treatment of modeled mixtures as replicate measurements expected to have no true differential expression from the mixture samples. Any detected differential expression between a mixture and its predicted expression values is indicative of a bias in the measurement process. In the BLM or SEQC datasets, differential expression was detected only in the ribosomal RNA genes (NR_003286.2, NR_003287.2, NR_023363.1). This detected differential expression reflects the sample to sample variance in rRNA depletion.

**Figure 6:**
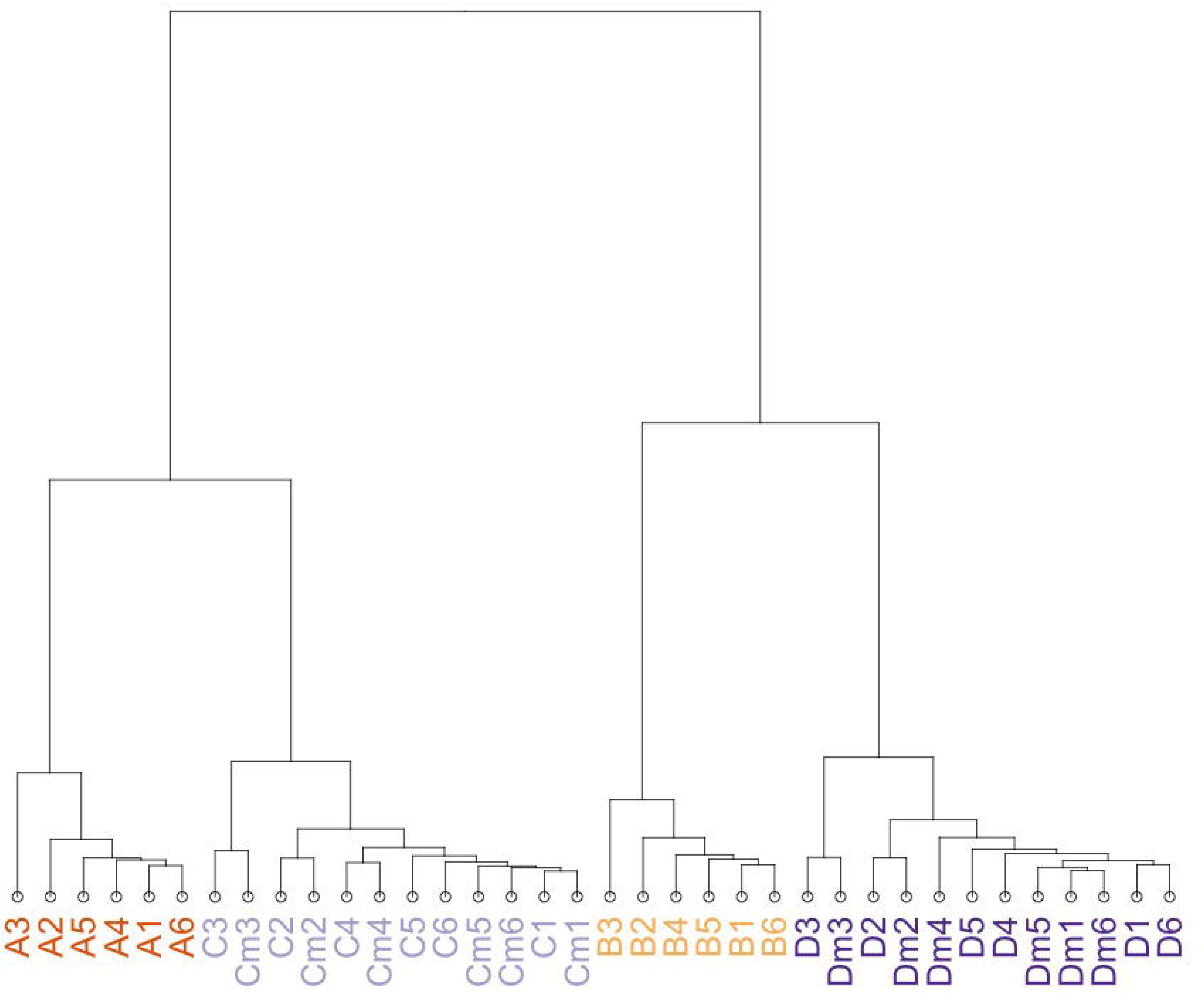
Clustering of Expression measures in 4 SEQC samples and 2 *in-silico* replicate samples across participating sites: The close agreement between modeled (Cm, Dm) counts and actual counts (A,B,C,D) at sites numbered 1-6 supports the validity of assumptions used to model Cm and Dm counts. Euclidian distance measures between samples show that the various samples are of greater distance from one another, while the *in-silico* modeled samples are most similar to the correct corresponding sample.

## Discussion

Mixtures of biological samples can be useful as process controls for measurements with linear response functions. A mixture can be treated as linear combination of its components. Two experimental datasets with known mixture parameters were used to test the linearity of RNA-seq measurements. In RNA-seq, the mRNA fraction of the total RNA mixture components must be accounted for in order to reflect the fact that mixtures of RNA are calculated based on mass fractions of total RNA and the sequencing experiments measures only mRNA.

Mixtures with either known or unknown proportions can be analyzed. If mixture proportion information is known *a priori*, genome-scale data can be used as a process control to test the repeatability and sensitivity of measurements by comparing observed and expected measures. Alternatively, if the mixture proportions are an unknown and desired parameter, expression measures from the mixture in combination with the single components can be used to experimentally determine the mixture proportions. This application can be valuable to un-mixing biological mixtures, including clinical mixtures, cell cultures, and xenografts[27-32]. While the mRNA fraction correction is required for RNA-sequencing measurements, the general mixture model is theoretically applicable to any measurement with a linear response function.

Mixtures can provide measurement process assurance to a sequencing experiment. Using mixture samples alongside pure samples, one can demonstrate the reproducibility and sensitivity of genome-scale RNA, protein, as well as metabolite measurements. The main goal of this type of mixture analysis is to create a known ratio value by which the measurement characteristics of an experiment can be assessed. While an experiment’s measurement of this known ratio is not sufficient to prove the validity of the measurement, it is a necessary condition, and any deviations are indicative of bias.

We demonstrate process control usage of mixtures by comparing the nine SEQC sites. Figure 5 shows a summary plot of the estimated component fractions for each sample. The dispersion and bias of the points from the target value give an indication of the overall process accuracy. Within this set of labs there are easily discernable changes, which could indicate process errors. Site 1 looks strong – there is no bias, and a modest and regular level of dispersion. In site 2 the dispersion of component C is exaggerated, suggestive of an issue in the handling of that particular sample. Site 4 has less dispersion than site 1, but has introduced a bias. Site 7 is from a completely different sequencing instrument, and shows that there is similar dispersion to the previous instrument, but a bit of a bias. However, site 8 shows that this bias does not occur in every run. This comparison of SEQC sites, shows that even these summary plots can detect differences between runs. It is for this reason that we suggest the use of mixtures as process controls for RNA-seq experiments. Comparing the dispersion and bias as you make changes to your experimental process allows you to evaluate the effect of these changes on the measurement quality. Text box 1 describes several types of changes that can be evaluated in this way.

While we demonstrate mixture analysis with two specific samples, the analysis is fully generalizable to any number or type of mixture components. Any mixture split into known individual components can be measured in this way. For example, a clinical researcher may have three samples of interest from healthy, chronically diseased and acutely diseased sources. A mixture of these three cell types would provide confidence in the measurements made on the three samples individually by verifying the repeatability of that measurement. It can also provide a benchmark sample to assess comparability over space and time. These mixtures can detect biases introduced by batch effects, operator effects, sample mislabeling, and technical artifacts while evaluating the variability of the measurement. Mixture samples with known proportions can help determine experimental reproducibility and discover technical artifacts introduced by the measurement process by comparison of the expected to observed proportions.

With this analytical model, end users and core facilities can use known mixtures as a process control to track changes in measurement quality whenever changes to the experimental process are made. By including a predefined mixture, cross-sample comparisons can be made to demonstrate the internal consistency of measurements made using any new experimental technique, kit, or downstream analysis tool. In this way, there is some assurance that changes in experimental protocol have not affected measurement reproducibility. Residuals from modeled counts can be used as a metric to evaluate the magnitude of effect an experimental process has on the linearity and precision of underlying measurements.

In addition to gaining an understanding of the measurement process using the benchmarking workflow, unknown samples can be collected and studied to determine the relative proportion of known components. Proportions of components can be determined even in the absence of any type-specific markers, given measurable differences in expression between the cell types.

Resolving the composition of mixtures has proven useful in determining the purity of cell lines or proportions of heterogeneous cells, in identifying interesting cellular contaminants such as partially differentiated cells, and understanding clinical samples containing mixed cell types. In contrast to approaches using transgene expression [41], the mixture model described here can evaluate tissue sample purity without focusing on a handful of tissue-specific genes, marker genes, or transgenes. We expect mixed-sample RNA to be useful in regulatory applications, where a demonstration that a therapeutic stem-cell mixture has a specific composition may be key to ensuring safety and efficacy [48].

### Spike-in controls measure mRNA content of samples

In addition to providing limit of detection and cross-experiment comparison characterizations of a dataset, spike-in controls can be used in mixture samples to determine the mRNA fraction of cells. mRNA fraction is a critical parameter for comparing samples that do not have identical total RNA content. This is most relevant to cells with variable global expression [24], including comparisons across and within cell cycle, tissues, and developmental states [40]. mRNA fraction is also critical in single cell gene expression studies, where lysis efficiency and total RNA content can vary greatly from cell to cell.

We demonstrate that the ERCC controls can be used as an estimator of mRNA content within samples. Of note, the SEQC study[34] results showed a large degree of variation in sample sequencing library preparation even at the same site, but that the sequencing library replicates prepared at a single site and then sequenced at multiple laboratories resulted in very consistent measurements between sites. Variation in library prep is primarily due to variability in mRNA enrichment, and is the primary source of variability in spike-in controls[39, 49].

There are many methods used to determine component gene expression profiles from mixture samples. To the best of our knowledge, our method is the only one that accounts for mRNA fraction. In RNA-seq experiments, mRNA fraction can be calculated with information obtained via spike-in controls. When comparing samples with variable mRNA content, bias arises when that variability is not accounted for. We describe a straightforward method for measuring the enrichment of mRNA in RNA-seq samples using spike-in RNA. We show that mRNA-corrected deconvolution of two mixture datasets returns the best approximation of known mixture proportions (Figure 4+5), demonstrating suitability for solving unknown mixtures of known components.

Previous methods used to determine the composition of RNA-seq mixtures make inaccurate estimates of mixture proportion in the BLM sample where the mRNA fractions vary substantially between mixture components. These methods are nearer to true values in the SEQC sample, where the mRNA fraction difference is less significant, but all estimates are improved by incorporating mRNA content (Supp. Figure 3).

### Recommendations for use

Control mixtures most easily demonstrate that an experimental process is linear and internally consistent, and can track the changes in variability over time. A first experiment with a new process should utilize these controls to demonstrate the reproducibility of measurements between single component and mixture samples. Subsequently, changes to the process can be evaluated by comparing the model residuals before and after the change. For example, a lab interested in changing from a total RNA measurement to a messenger RNA measurement may wish to evaluate if this change had any effect on sequencing output. The change in the sum of residuals between these two different experiments would allow a global comparison, while the change in residuals of individual genes may highlight a set of genes, which become inconsistently biased between experiments. Text box 1 shows three potential use cases for mixtures used as process control.

| User | Sample Design | Time to use | Benefit |
| --- | --- | --- | --- |
| Technology and Experimental Protocol | Mix 2-3 components of biological interest | When validating experimental protocols and/or | Demonstrate the ability of a protocol and/or technology |
| **Developers** | appropriate for use to evaluate the experimental protocol/technology | technology platforms | platform to consistently measure transcripts of interest |
| **Core Labs** | Repeated measures of highly-available samples | After changes in technician, protocol, reagents, or technology platform, etc. | Show internal consistency of output and linearity of measurements<br>Identify biases<br>Understand dispersion |
| **Informaticians** | Relevant samples, public data (eg: SEQC, this study) | When comparing informatics analysis tools or developing new tools | Use non-simulated benchmark datasets to determine how accurate results are. |

### Limitations

Although mean mixture proportion values returned from a linear combination of mixture components approximate the nominal mixture proportion in both measured samples, the increased variability of the muscle estimate in the BLM mixture (error bars, Figure 4) suggests that there is a lower limit to being able to determine low-abundance mixture components. Due to mRNA fraction, the muscle component of the BLM mix was as low as 10 percent of sequenced RNA in BLM-2. It may be possible to determine lower-proportion mixture components with confidence, but this study did not generate the required data to do so.

Our estimation of mRNA fraction is imperfect; an assumption of the model we built is that the mRNA fraction is constant between replicates of the same sample. Supplemental Table 1 shows that the mRNA fraction varies by as much as 5 percent from library to library. This variability is a source of error in our model. The variability in mRNA fraction is likely due to batch effects in the mRNA enrichment process. This hypothesis is reinforced by the prevalence of non-mRNA transcripts incorrectly called as differentially expressed between mixture replicates. The sequencing technology and library preparation methods used in these experiments also added limitations to the experiments. These are described in supplemental note 1.

## Conclusions

We demonstrate the linear response function and specificity of RNA-sequencing measurements using mixtures of biological samples. We recommend the use of such mixtures as benchmarks to characterize the repeatability and reproducibility of experiments. Spike-in controls can be used to calculate the mRNA content of total RNA mixtures, compensating for biases introduced by mRNA enrichment. Our method creates a framework for using mixtures in measurement process control and corrects for biases introduced by ribosomal depletion. Using an mRNA fraction correction improves the accuracy of mixture proportion determination in RNA-seq experiments.

Benchmarking genome-scale measurements using mixed samples will remain useful even after the era of short-read sequencing is over. Answering the biological question of “what types of cells are in the mixture I’m sequencing?” requires more information than even a perfect transcriptome reconstruction could provide. The biological and measurement value added by mixed samples are demonstrated here to be platform-independent. We anticipate that mixtures can provide the same measurement assurance to protein and metabolite measurements. Confidence in the reproducibility of measurement and understanding the components in complex biological samples will always be a staple of quality science.

### Methods

#### Library Preparation

For the BLM experiment, Human Brain Reference RNA, Human Liver Total RNA, and Human Skeletal Muscle Total RNA were purchased from Ambion. This purified RNA was quantified by absorbance on a NanoDrop 1000, mixed in the specified proportions, then spiked with ERCC RNA transcribed from NIST SRM 2374. For Illumina sequencing, the Illumina TruSeq protocol was followed. HiSeq runs generated 100+100bp paired-end reads. Solid 5500 sequencing followed the Life Technologies Whole Transcriptome protocol, yielding 75+35 bp paired-end reads. Spike-in composition and amounts are included in the data submission to ENA.

#### Quantitation and Data Normalization

BLM gene counts were based on raw count data quantified using HTSeqCounts [40]based on a variety of genome and transcriptome references [42-45] after mapping reads to the genome with Topha t[46]. Raw counts were then normalized using the upper quartile method implemented in EdgeR [36]. Supplemental Figure 3 utilizes RSEM [47]. HTSeq-counts version 0.5.4 was run with options to deal with non-stranded reads in the intersection-nonempty mode. The SEQC data used are available as count tables from GEO GSE47774.

#### Calculating Unknown Mixture Estimates

The relative abundance of components in unknown mixtures were calculated by first observing the mean mRNA fraction for the neat components across replicates. The count data in the mixture was set as the response, predicted by the count data from the individual components modified by the mRNA fraction, as based on the mixture equations. An example R script ‘generalmixturesolver’ is provided at http://github.com/usnistgov/mixtureprocesscontrol as a supplemental file to clarify this procedure.

#### Availability of supporting data

The SEQC data is available from GEO GSE47774.

[http://www.ncbi.nlm.nih.gov/geo/query/acc.cgi?acc=GSE47774]

The BLM data is available from the European Nucleotide Archive, PRJEB8231.

[http://www.ebi.ac.uk/ena/data/view/PRJEB8231]

Figure code, count tables, and example scripts available on https://github.com/usnistgov/mixtureprocesscontrol

List of Abbreviations: ERCC - External RNA Control Consortium, TPM – Transcripts per Million, FPKM – Fragments per Kilobase per million mapped reads, mRNA – messenger RNA

#### Competing Interests

The authors declare that they have no competing interests.

Certain equipment and instruments or materials are identified in the paper to adequately specify the experimental details. Such identification does not imply recommendation by the National Institute of Standards and Technology, nor does it imply the materials are necessarily the best available for the purpose.

#### Author Contributions

Analysis by JP, MS, PP, MM, and SM. Manuscript by JP. Experimental design by PP, SM, JM, MS, and MM. Sample preparation and sequencing by JM+NCI sequencing core.

## Acknowledgements

The authors would like to thank Dr. Steve Lund for helpful feedback around this work.

## Supplemental Figures

**Supplemental Figure 1:**
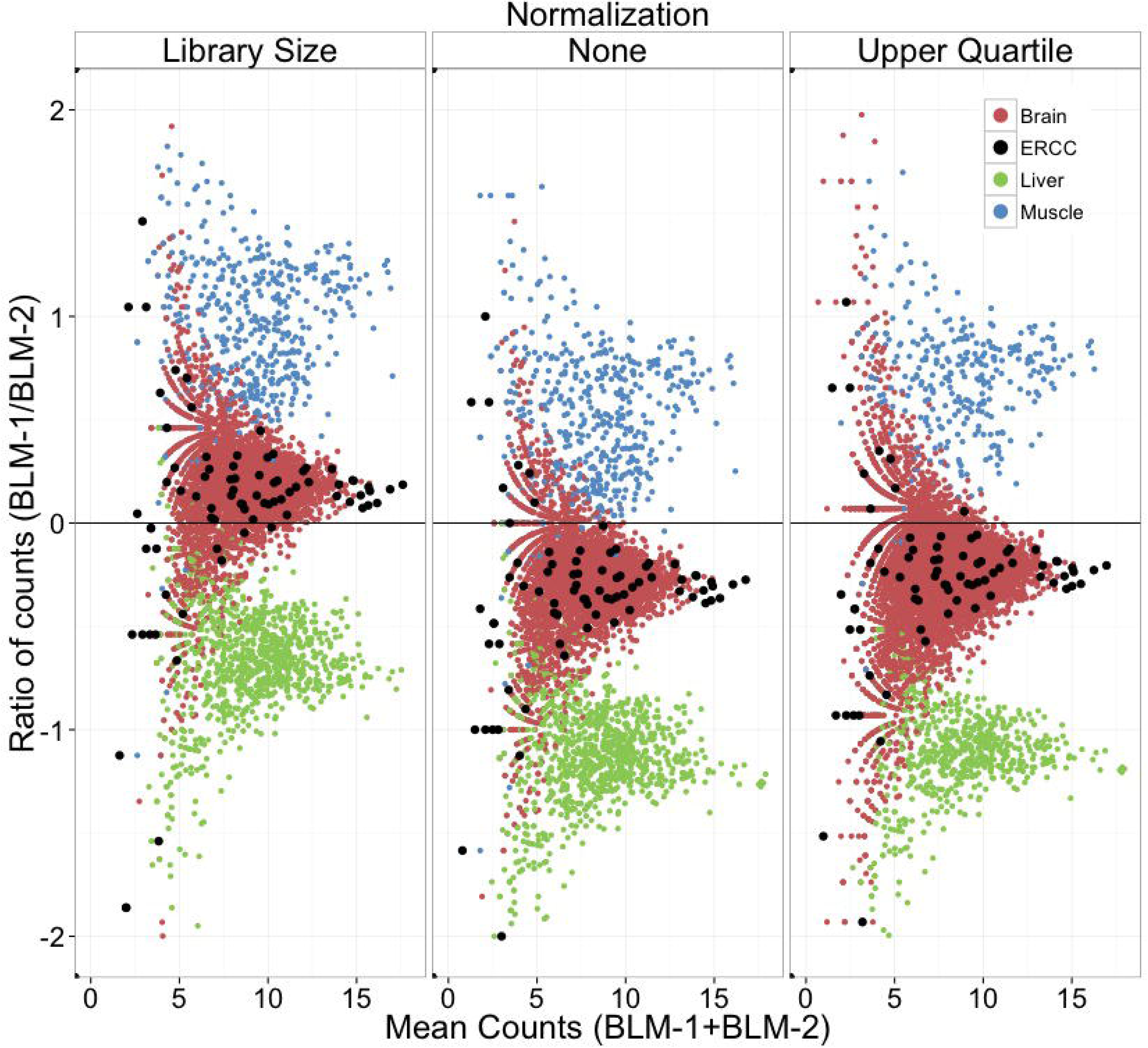
Bland-Altman log-ratio(M) - log average(A) plots comparing gene expression in BLM-1 to BLM-2, which were mixed with a designed ratio of 1:1 brain RNA, 2:1 muscle RNA and 1:2 liver RNA. Points representing gene expression values for genes expressed at 5-fold greater levels in a specific tissue are colored based on the tissue in which they are selectively expressed. Non-tissue selective mRNAs are omitted for clarity. Library size normalization scales all libraries to a common total number of counts, while upper quartile normalization scales to the 75^th^ percentile of the counts for each library. None of these normalizations accurately reflects the designed ratio of transcripts between samples.

**Supplemental Table 1:** Message RNA fraction (*ρ*) calculations as a function of spike amount. Spike mass is accounted for in the mRNA fraction calculation. The spike-ins varied by amount (“u” or “d” samples) and content (pools ‘a’ or ‘b’) in both tissue mixtures (1 and. 2). Calculated mRNA fractions vary by +/- 6% across these 10 BLM mixtures, showing that the calculation is robust to spike-in mass and content. mRNA fraction calculations for the ERCC pools must account for the 3-plex nature of the mixes. The shown ratios are for the subset of spike-ins which are present at a 1:1 ratio in each sample.

|  | BLM1-a | BLM1-ad | BLM1-au | BLM1-b | BLM1-bd | BLM1-bu | BLM2-a | BLM2-b | BLM2-bd | BLM2-bu |
| --- | --- | --- | --- | --- | --- | --- | --- | --- | --- | --- |
| Count<br>Ratio | .0695 | .0095 | .6698 | .0719 | .0098 | .6342 | .0706 | .0737 | .0098 | .6649 |
| Spike<br>Added | .08 | .01 | .64 | .08 | .01 | .64 | .08 | .08 | .01 | .64 |
| message<br>fraction $\rho$ | 1.152 | 1.058 | .955 | 1.112 | 1.017 | 1.009 | 1.132 | 1.085 | 1.025 | .962 |

**Supplemental Figure 2:**
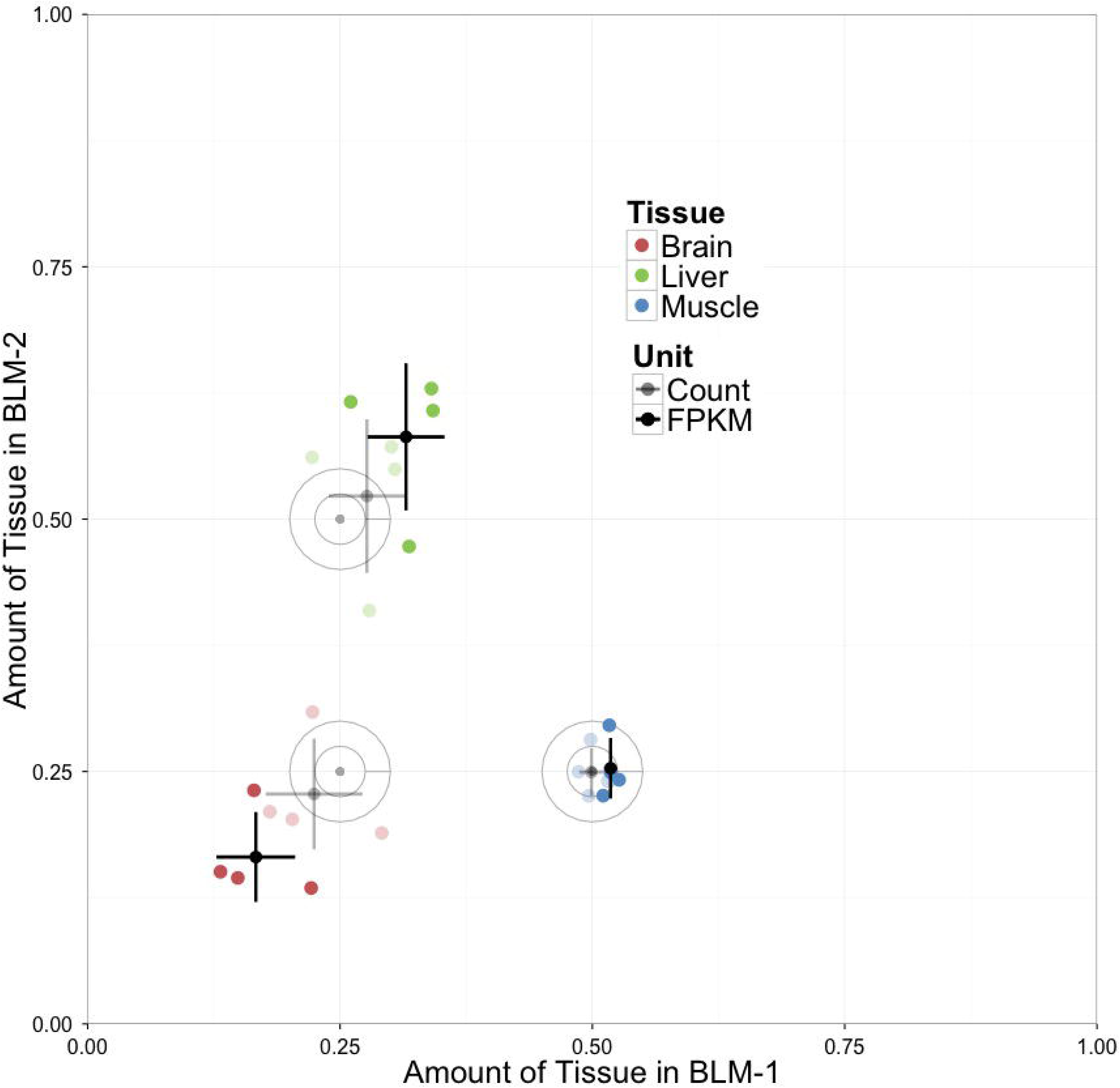
The effect of using FPKM units. Estimates of mRNA fraction (light points are calculated using count values, dark points using FPKM values) result in a relatively poor solution to the mixture proportion. Both data types are taken from the same RSEM output.

**Supplemental Figure 3:**
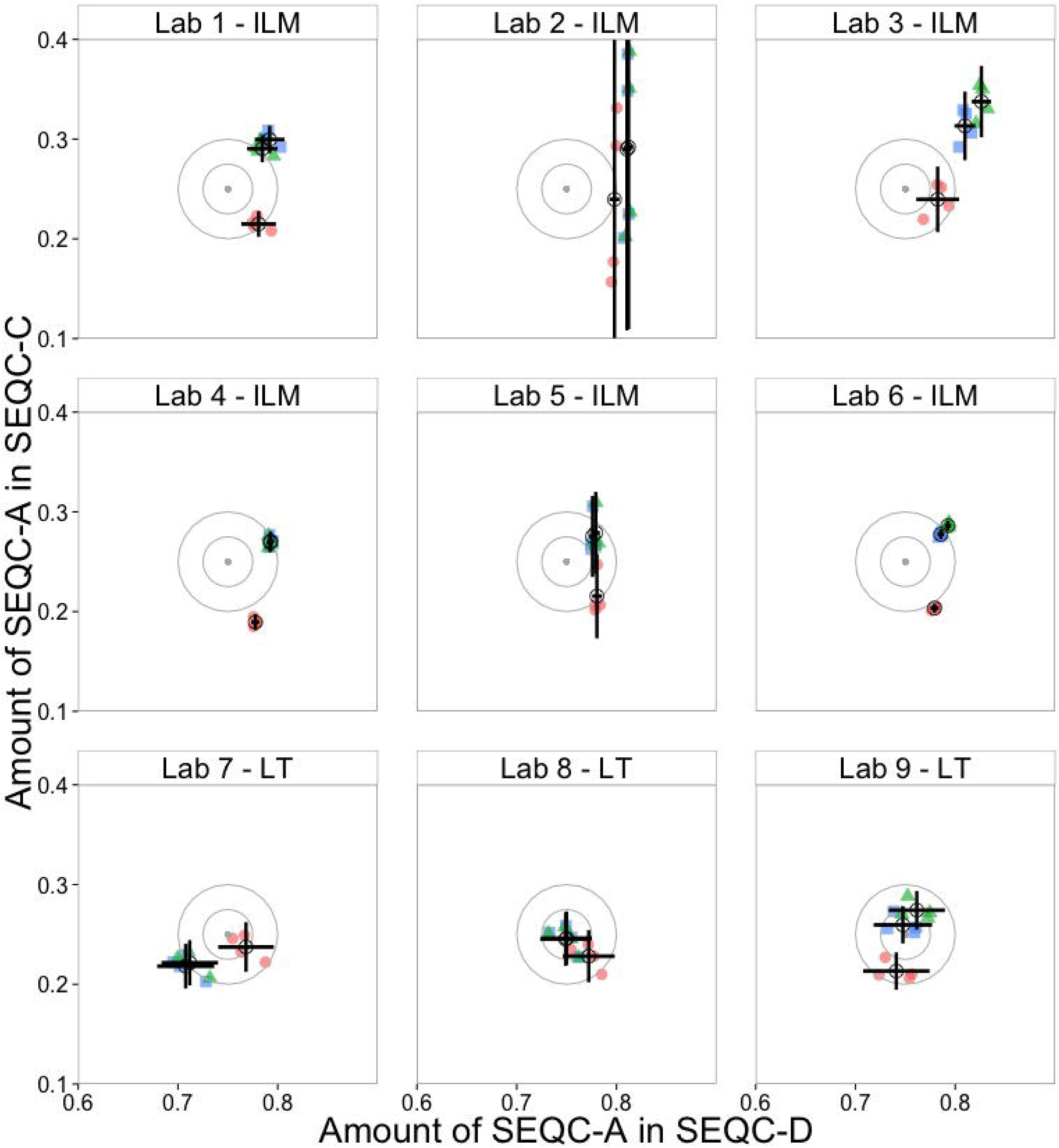
Mixture proportions returned by a simple model (Equation 1, blue circles), by an mRNA-corrected model(*ρ-corrected* mixture equations, green triangles) and by the DeconRNASeq package[36] (red diamonds) on SEQC data. Lab # - LT and - ILM indicate the manufacturer of the sequencer used at each participating lab (Life Technologies and Illumina, respectively).DeconRNASeq implements the same general idea, but lacks mRNA fraction correction. In the SEQC data, there is a relatively small mRNA fraction difference between samples, but significant improvements are achieved by correcting for the mRNA fraction. The mean distance from true value across all labs is 0.052(Simple model), 0.033(mRNA-corrected), and 0.048(DeconRNASeq). Error bars represent the SD of four independent libraries from the same RNA source.

## Supplemental Note 1

RNA-seq is capable of making transcript isoform-specific measurements. However, long reads of high depth are required to adequately differentiate between isoforms. Investigations of isoform-level measurements from the BLM dataset, (Table 2) which utilized 75x35bp paired-end reads on the 5500 and 100x100bp paired-end reads on the HiSeq, showed that while the model is extensible towards such measurements, the reduced mean read counts make transcript isoform-level expression measurements less precise due to shorter read length and lower sequencing depth. 92 percent of genes were modeled to within 1 log2 unit of the measured value, while only 85 percent of transcripts were.

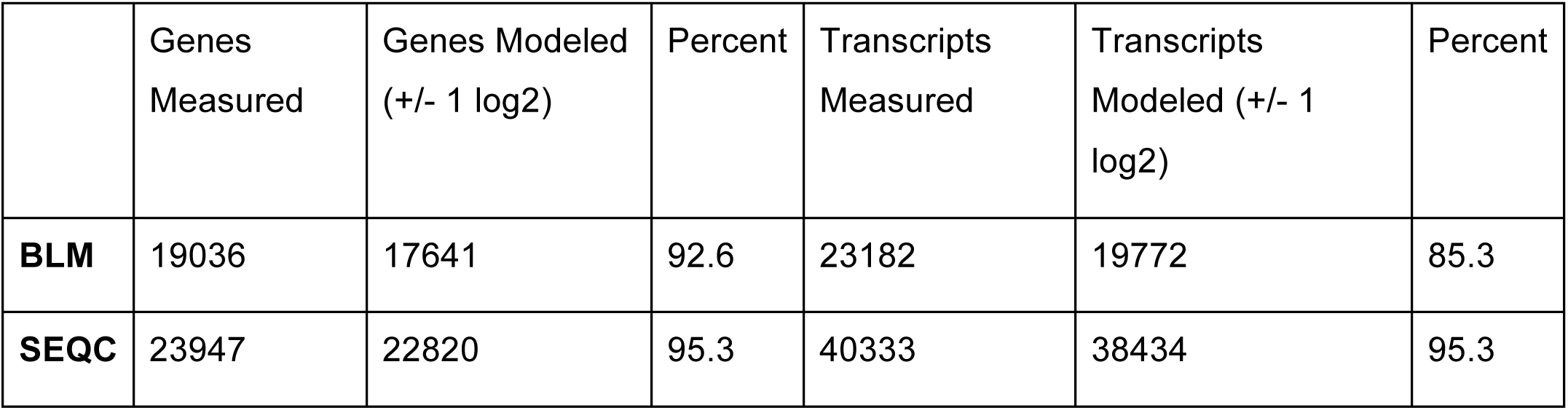

The substantially increased read depth in the SEQC experiment led to 95% of both isoforms and genes being consistently modeled. In the SEQC dataset, 95% of detected isoforms could be consistently modeled to within a factor of 2, and the same percentage of genes could be reasonably predicted. After applying a variance-stabilizing transformation using DEseq[38], every gene and transcript (100%) in the SEQC dataset were correctly modeled by these criteria. The BLM dataset does not contain sufficient replication for variance-stabilizing analysis.

